# Monitoring insecticide resistance in mosquitoes from Tunisia: implications for vector control strategies

**DOI:** 10.64898/2026.05.21.726919

**Authors:** Latifa Remadi, Natasa Kampouraki, Emmanouil Alexandros Fotakis, Linda Grigoraki, Sofia Balaska, Samia Layouni, Hamouda babba, Najoua Haouas, John Vontas

## Abstract

Tunisia is considered a high-risk country for arboviral transmission, and the re-emergence of malaria. Vector control programs predominantly rely on insecticide-based interventions, however knowledge on the insecticide resistance status of local vectors is limited. Here, we determined the mosquito species composition and investigated the resistance status of disease vectors to insecticides, collected from two distinct ecological settings; urban setting and rural setting. Known insecticide resistance loci were molecularly analysed. A total of 210 mosquitoes were collected, representing three main genera: *Culex* (n = 119), *Aedes* (n = 79), and *Anopheles* (n = 1). Within the *Culex* genus, *Cx. pipiens pipiens* and *Cx. pipiens* hybrids were the predominant species in Monastir and Kairouan respectively. Amongst *Aedes, Ae. albopictus* was the most abundant species in Monastir, while *Ae. caspius* predominated in Kairouan. The knockdown (*kdr*) mutation L1014F was detected at both sites across the three *Cx. pipiens* biotypes. In contrast, *Cx. perexiguus* showed no *kdr* mutation in Kairouan and a low frequency in Monastir. Among *Aedes* species, *kdr* mutations were detected exclusively in *Ae. albopictus*, restricted to domain III (mutation I1532T). No mutations in the *chs* gene were identified in both populations. *Aedes albopictus* collected in Monastir exhibited a moderate copy number variation for both carboxylesterases 3 and 6. No mutations were detected in the *Anopheles labranchia*. Our study results indicate the suitability of diflubenzuron for *Culex* and *Ae. albopictus* control in central Tunisia, emerging pyrethroid resistance in both vector species, and possibly reduced organophosphate efficacy against *Ae. albopictus*.

## Introduction

Tunisia, located in North Africa on the Mediterranean coast, occupies a strategic biogeographical position bridging Europe and Africa. The country’s diverse ecosystems, spanning from humid coastal plains to arid inland regions support a rich mosquito fauna including several *Anopheles, Culex*, and *Aedes* vector species (Ahmed Tabbabi, 2017), favouring the emergence and transmission of vector-borne diseases..

Notably, Tunisia was an ancient site for malaria in North Africa. Although the disease was eradicated in 1977 (Chadli et al., 1986) the risk of its re-establishment remains significant, driven by the presence of suitable mosquito breeding sites, documented vector presence such as *An. labranchiae, An. claviger sensu stricto, An sergenti* and *An. multicolor*, favourable environmental conditions, and the continuous arrival of potentially infected individuals from sub-Saharan Africa (Ouni et al., 2026).

*Culex pipiens*, the main vector of West Nile virus (WNV) is ubiquitous across Tunisia and plays a key role in nuisance biting and disease transmission, with sympatric occurrence of *Cx. pipiens molestus, Cx. pipiens pipiens* and their hybrids (Beji et al., 2017). WNV is considered a well-established public health concern in Tunisia, with several episodes of human disease reported over the past decades. The country experienced two major WNV outbreaks, first in 1997 and later in 2003 (Triki et al., 2001, Hachfi et al., 2010, Riabi et al., 2014), which highlighted the virus’s capacity to re-emerge and spread locally. Evidence of WNV circulation has since been documented in both equine populations and wild birds, confirming the persistence of the virus in natural hosts and reservoirs (Ben Hassine et al., 2014, Hammouda et al., 2015). In addition, WNV lineage 1 has been detected in *Cx. pipiens* collected from central Tunisia, underscoring the important role played by this widespread species in local transmission cycles (Wasfi et al., 2016).

The recent detection of *Aedes albopictus* in Tunisia has raised new concerns (Bouattour et al., 2019). As a highly adaptable invasive mosquito and a competent vector of dengue, chikungunya, and Zika viruses, its establishment in urban and peri-urban settings introduces an additional layer of risk for future arboviral outbreaks.

The prevention and control of mosquito-borne diseases, as well as the alleviation of nuisance caused by mosquitoes strongly depends on the implementation of vector control programs that utilize chemical, mechanical, and biological interventions.

In Mediterranean countries, vector control programs have traditionally relied on chemical interventions, mainly using insecticides such as pyrethroids, carbamates (CRB), and insect growth regulators (e.g., diflubenzuron (DFB)), while organophosphates (OP) were historically used but are now increasingly restricted or phased out in many regions (Cholvi et al., 2026).

In Tunisia chemical control has long been at the heart of mosquito management, starting from the malaria eradication era between 1966 and 1972 (Chadli et al., 1986) involving the intensive use of organochlorines, OPs, CRBs, and pyrethroids (Tabbabi & Daaboub, 2018a, Tabbabi & Daaboub, 2018b). More recently, mosquito control relies primarily on larval source management, with organophosphate larvicides such as temephos historically applied against *Culex pipiens* larvae. Additionally, biological larvicides like *Bacillus thuringiensis* var. israelensis, are increasingly considered as safer alternatives with lower non-target impacts (Zribi Zghal et al., 2018). In emergency situations (e.g., high vector densities or outbreak risk), control programs may incorporate pyrethroid-based adulticide and enhanced source reduction within integrated vector management frame works (Tabbabi & Daaboub, 2018b).

Although biocides are necessary for vector borne diseases (VBD) prevention and control, a serious global threat associated with their extensive use is the development of insecticide resistance in vector species, undermining vector control efforts. Insecticide resistance predominantly arises through two main mechanisms: toxicodynamic mechanisms which involve mutations at insecticide target sites reducing the binding efficiency of the compound; and toxicokinetic mechanisms which limit the effective dose of insecticide within the insect through enhanced metabolic detoxification and other pathways (i.e. sequestration, reduced cuticular penetration, increased excretion) (Liu, 2015).

Well studied target-site mutations include knockdown resistance (*kdr*) mutations in the voltage-gated sodium channel (*vgsc*), associated with pyrethroid resistance in several *Anopheles* species (L1014F/C/S), *Culex pipiens* (L1014F/C/S) and *Aedes albopictus* (F1534C/L/S, V1016G/I, and I1532T). Other well studied cases are the mutations F290V and G119S on acetylcholinesterase (*Ace*-1) associated with resistance to OPs and CRBs in *Culex pipiens* and the mutations in chitin synthase (*chs-1*) associated with resistance to the larvicide DFB in *Culex pipiens* (I1043L/M/F)(Paronyan et al., 2024).

Metabolic resistance involves the overexpression of detoxification enzymes (e.g cytochrome P450 monooxygenases (P450s), glutathione S-transferases (GSTs), and carboxylesterases (CCEs)) such as CCEae3a and CCEae6a, which have been associated with temephos resistance in *Aedes albopictus* (Grigoraki et al., 2017).

In Tunisia, variants at both esterase genes and the ace locus of *Cx. pipiens* populations have been detected, indicating OP resistance and long-term selection pressure (Ben Cheikh et al., 2009). More recently, bioassays of adult *Cx. pipiens pipiens* from three Tunisian districts demonstrated resistance to the pyrethroid deltamethrin (Tabbabi et al., 2018).

Among malaria vectors, resistance to permethrin and deltamethrin has been documented in *An. sergentii* from southern Tunisia (Tabbabi & Daaboub, 2018b). For *An. labranchiae*, a study found exceedingly high resistance ratios to temephos in some field populations (RR□□ up to 624), levels not previously reported in North Africa, highlighting the urgent need for molecular and biochemical investigations (Tabbabi & Daaboub, 2018a).

Despite these first reports documenting insecticide resistance in important vectors, studies addressing insecticide resistance in Tunisian mosquito populations remain limited, leaving a critical knowledge gap for vector control programmes. Additionally, to date, there is no information regarding insecticide resistance status in *Ae. albopictus* (Bouattour et al., 2019) (Bohers et al., 2020).

Our study aims to address these knowledge gaps by performing molecular analyses to elucidate the mechanisms of resistance in mosquitoes collected from urban and rural zones in Tunisia.

## Materials and Methods

### Study regions and field sample collections

Mosquitoes were collected in two contrasting sites in Tunisia: an urban area in Monastir and a rural in Kairouan. Monastir is a highly touristic coastal city located in the central eastern region of Tunisia. The presence of coastal saltpans in Monastir supports migrating and wintering birds, whilst agriculture is less dominant and less intensively irrigated compared to major farming zones. Kairouan lies inland and represents a major agricultural area, where irrigated crops and livestock farming are highly developed. Sampling took place between August and September over two consecutive years (2024–2025). Every week, three Center for Disease Control (CDC) light traps were set up near animal shelters at sunset and retrieved at sunrise. Prior to each collection, landowners were contacted, and their consent was obtained for trap placement. Collected mosquitoes were sorted by species and preserved in 70% ethanol at −20 °C until molecular analyses were performed.

### Molecular identification of mosquito species

DNA was extracted from each individual mosquito using DNAZOL, following the protocol provided by the manufacturer’s instructions (Invitrogen, Carlsbad, CA, USA). DNA purity and quantity was assessed using Nanodrop 2000c spectrophotometer (Thermo Scientific). Molecular identification was performed based on the mitochondrial cytochrome oxidase I gene (COI) followed by sequencing as described in (Folmer et al., 1994). Obtained sequences were analysed and species were identified with BLASTn (http://blast.ncbi.nlm.nih.gov/) based on homology with available mosquito sequences deposited in GenBank. *Culex pipiens* were identified based on polymorphisms in the intron region of the *Ace*-2 gene. The species biotypes (pipiens, molestus and hybrid) were discriminated using PCR amplification of the 5’ flanking regions of the microsatellite locus CQ11 (Bahnck & Fonseca, 2006).

### Insecticide resistance monitoring

*Culex* individuals were genotyped for the pyrethroid-associated *kdr* mutations (L1014F/C/S) in the *vgsc* gene. Resistance to OP and CRB was examined through detection of the *Ace*-1 G119S and F290V mutations using PCR–RFLP (Weill et al., 2004, Alout et al., 2007). We additionally investigated the I1043L/M/F substitutions in the *chs*-1 gene, which are linked to resistance to DFB (Paronyan et al., 2024). For *Aedes*, pyrethroid resistance was assessed by screening for six major *vgsc kdr* variants, (i.e. V1010G, V1010GI, I1532T, F1534C, F1534L and F1534S) (Paronyan et al., 2024). Moreover, the copy-number variation (CNV) of the CCEae3a and CCEae6a was also assessed. CNV levels were measured using qPCR following (Balaska et al., 2020) and (Grigoraki et al., 2017), with histone 3 and ribosomal protein L34 serving as reference genes. For Anopheles, both the *kdr* (L1014F/C/S) and the *Ace*-1 mutations were screened (Djadid et al., 2009, Weill et al., 2004).

## Results

### Mosquito fauna composition

A total of 210 mosquitoes were collected from the two study sites, Monastir (N=120) and Kairouan (N=90), belonging to three genera: *Culex, Aedes, Culiseta* and *Anopheles*. In Monastir, the dominant species was *Ae. albopictus*, followed by *Cx. perexiguus, Cx. pipiens* and *Ae. caspius* (Table 1). Other species such as *Cx. laticinctus, Culiseta longiareolata*, and *An. labranchiae* were also recorded, though in low numbers. In contrast, mosquito fauna in Kairouan was dominated by *Cx. perexiguus, Cx. pipiens*, and *Ae. caspius. Culex theileri* was found in a low number, while no *Ae. albopictus* were detected in this rural site. For *Cx. pipiens* biotype, the dominant species in Monastir was *Cx. pipiens pipiens* (56%) followed by *Cx. pipiens hybrid* (24%) and *Cx. pipiens molestus* (20%). In Kairouan, the dominant biotype was *Cx. pipiens hybrid* (46%) followed by *Cx. pipiens molestus* (38.5%) and *Cx. pipiens pipiens* (15.5%).

**Table 1.**
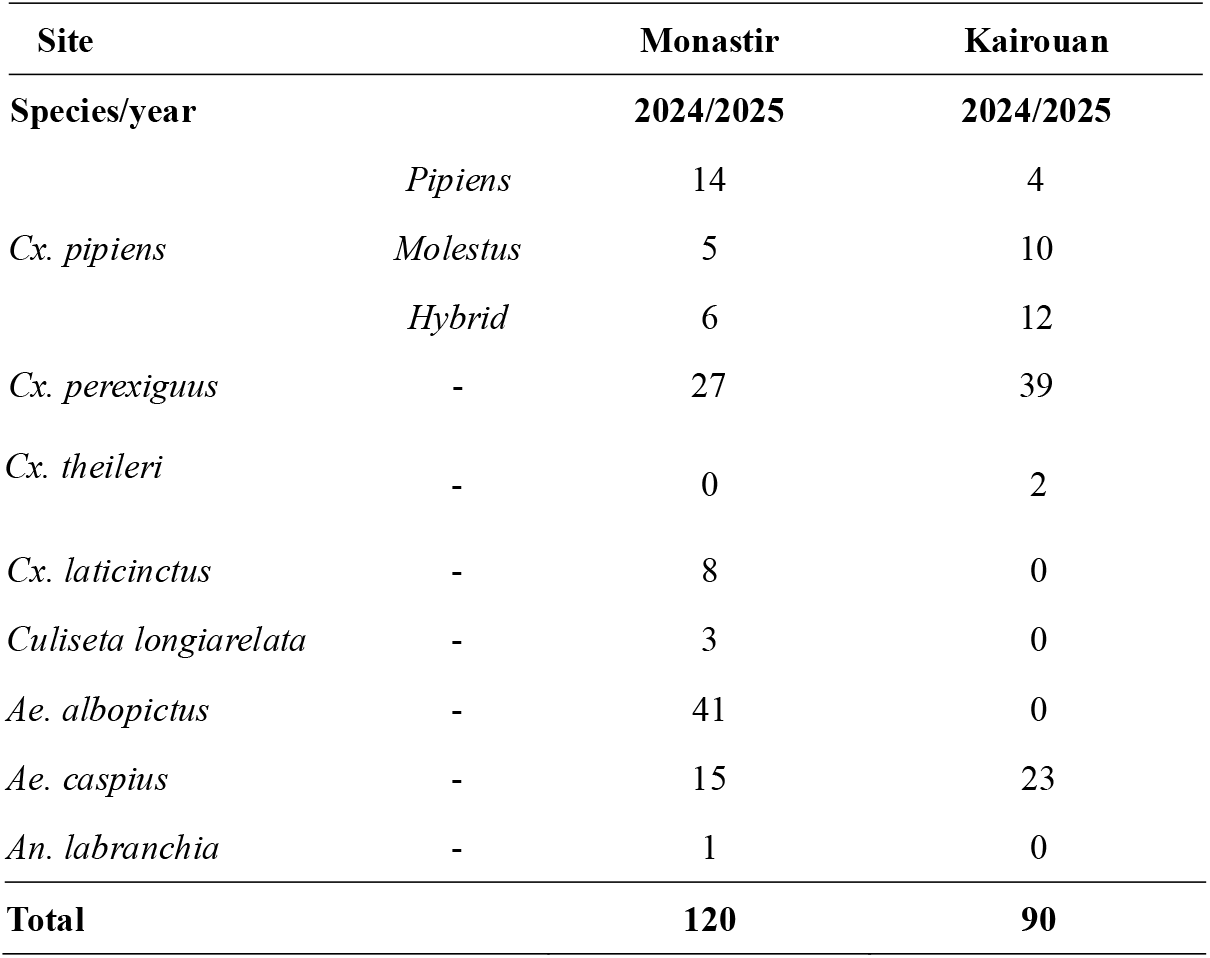
Mosquito fauna composition.

### Molecular Detection of Resistance Markers

#### Genotyping of target site mutations in *Culex* species

Three well known insecticide resistance markers were screened for in *Cx. pipiens* and *Cx. perexiguus* populations from Monastir and Kairouan: the *kdr* L1014F mutation in the *vgsc*, the *Ace-1* G119S and F290V mutations, and the *chs* I1043L/M/F mutations. Mutation L1014F was detected in both sites and in the three *Cx. pipiens* biotypes, with higher allelic frequencies in *Cx. pipiens hybrid* in Kairouan (36.36%) and *Cx. pipiens molestus* (75%) in Monastir (Table 2). Overall, heterozygous individuals (L/F) were more frequent than homozygotes for the resistance mutation (F/F) (Table 2). Homozygotes for the resistance allele were more frequent in Monastir (52.1%) compared to Kairouan (9.1%). For *Cx. perexiguus*, the *kdr* mutation was nearly absent, detected only in a single specimen from Monastir with a low frequency (4.1%). *Ace*-1 mutation G119S was present at a low frequency in *Cx. pipiens* from both sites, albeit was not detected in *Cx. perexiguus*. Mutation F290V was not detected in any *Culex* samples. Similarly, we didn’t detect any *chs* I1043L/M/F mutations, in *Culex* mosquitoes.

**Table 2.** Monitoring insecticide resistance in *Culex* species.

| Mutation |  | *kdr*/ L1014F |  |  |  |  |  |  |  |
| --- | --- | --- | --- | --- | --- | --- | --- | --- | --- |
| Site | Species | N | SS | RS | RR | SS (%) | RS (%) | RR (%) | R alleles (%) |
| Kairouan | *Cx. perexigus* | 6 | 6 | 0 | 0 | 100 | 0 | 0 | 0 |
|  | *Cx. pipiens pipiens* | 8 | 6 | 2 | 0 | 75 | 25 | 0 | 25 |
|  | *Cx. pipiens molestus* | 14 | 12 | 2 | 0 | 85.7 | 14.28 | 0 | 14.28 |
|  | *Cx. pipiens hybrid* | 22 | 16 | 4 | 2 | 72.7 | 18.2 | 9.1 | 36.36 |
| Monastir | *Cx. perexigus* | 13 | 12 | 1 | 0 | 92.3 | 7.6 | 0 | 7.6 |
|  | *Cx. pipiens pipiens* | 28 | 17 | 9 | 2 | 60.7 | 32.1 | 7.1 | 46.42 |
|  | *Cx. pipiens molestus* | 8 | 4 | 2 | 2 | 50 | 25 | 25 | 75 |
|  | *Cx. pipiens hybrid* | 10 | 6 | 2 | 2 | 60 | 20 | 20 | 60 |

| Mutation |  | *Ace-1*/ G119S |  |  |  |  |  |  |  |
| --- | --- | --- | --- | --- | --- | --- | --- | --- | --- |
| Site | Species | N | SS | RS | RR | SS (%) | RS (%) | RR (%) | R alleles (%) |
| Kairouan | *Cx. perexigus* | 7 | 7 | 0 | 0 | 100 | 0 | 0 | 0 |
|  | *Cx. pipiens pipiens* | 8 | 7 | 1 | 0 | 87.5 | 12.5 | 0 | 12.5 |
|  | *Cx. pipiens molestus* | 16 | 15 | 1 | 0 | 93.7 | 6.25 | 0 | 6.25 |
|  | *Cx. pipiens hybrid* | 22 | 19 | 3 | 0 | 86.4 | 13.6 | 0 | 13.63 |
| Monastir | *Cx. perexigus* | 13 | 13 | 0 | 0 | 100 | 0 | 0 | 0 |
|  | *Cx. pipiens pipiens* | 26 | 24 | 2 | 0 | 92.3 | 7.7 | 0 | 7.6 |
|  | *Cx. pipiens molestus* | 8 | 7 | 1 | 0 | 87.5 | 12.5 | 0 | 12.5 |
|  | *Cx. pipiens hybrid* | 10 | 9 | 1 | 0 | 90 | 10 | 0 | 10 |

#### Genotyping of target site mutations in *Aedes* species

Two *kdr* loci in domain II (V1016G) and domain III (I1532T and F1534C/L/S) of the *vgsc* and the *chs* I1043L/M/F mutation were analysed in *Ae. albopictus* and *Ae. caspius* populations. In *Ae. albopictus* from Monastir, the V1016G mutation was absent, while mutation I1532T was detected at a low frequency (8.9%) (Table 3), and solely in the heterozygote state. No *chs* mutations were identified in this population. No *kdr* or *chs* mutations were detected in the *Ae. caspius* populations from Monastir and Kairouan.

**Table 3.** Monitoring insecticide resistance in *Aedes* species.

| Mutation |  | *kdr* III/ I1532T |  |  |  |  |
| --- | --- | --- | --- | --- | --- | --- |
| Site | Species | N | SS | RS | RR | R alleles (%) |
| Kairouan | *Ae. caspius* | 4 | 100 | 0 | 0 | 0 |
| Monastir | *Ae. albopictus* | 78 | 71 | 7 | 0 | 8.9 |
|  | *Ae. caspius* | 26 | 100 | 0 | 0 | 0 |

#### Detection of esterase gene amplification for *Ae. albopictus*

Of the 41 *Ae. albopictus* collected in Monastir, 14 individuals were screened for CNV in the *CCEae3a* and *CCEae6a* genes. Amplification of the esterase genes was detected. Most samples exhibited a moderate CNV, ranging from 2–16 copies for *CCEae3a* and 2–15 copies for *CCEae6a*. Overall, 12 out of the 14 screened mosquitoes (85.7%) displayed a moderate CNV for both *CCEae3a* and *CCEae6a* (Fig 1).

**Fig 1.** Copy number variation of *CCEae3a* and *CCEae6a* in *Ae. albopictus*.

#### Genotyping of target site mutations for *Anopheles* species

Only one *An. labranchia* specimen was identified in Monastir, and no insecticide resistance mutations (L1014F/C/S) were detected.

## Discussion

This study provides the first comprehensive molecular assessment of insecticide resistance in mosquito populations from central Tunisia. Here we report the mosquito fauna composition in Monastir and Kairouan, and the presence of *kdr* mutations in *Cx. pipiens* and *Ae. albopictus*, the *Ace-1* mutation in *Cx. pipiens* and a moderate CNV of *CCEae3a* and *CCEae6a* in *Ae. albopictus*.

Monastir, a coastal urban and touristic area with limited agricultural activity and irrigation, was dominated by *Ae. albopictus* and *Cx. pipiens* followed by *Cx. perexiguus*. Whereas Kairouan, an agriculture rural region with extensive irrigated crops and livestock, supported higher numbers of *Ae. caspius, Cx. perexiguus* and *Cx. pipiens*. These differences likely reflect variation in breeding site availability and human-mediated environmental changes. This distribution highlights the ecological contrast between the coastal urban environment and the rural irrigated landscape of Kairouan. Only a single *An. labranchiae* was collected in Monastir, which likely reflects the limited attractiveness of CDC light traps for this species. Previous studies have shown that human-baited double net traps are more effective in attracting *Anopheles* mosquitoes than CDC light traps (Degefa et al., 2020).

Aedes albopictus was first reported in northern Tunisia in 2018 (Bouattour et al., 2019), and the present study represents the first record of this species in central Tunisia. This finding indicates the expansion of *Ae*.□*albopictus* toward the center of the country likely driven by the specie’s ecological plasticity and human-mediated transport (Battaglia et al., 2022). Moreover, the absence of *Ae. albopictus* in Kairouan does not necessarily indicate that the region is free of this species, particularly because CDC light traps were used for sampling, which may be less attractive to this mosquito. Therefore, additional surveillance efforts specifically targeting *Ae. albopictus* are needed to better characterize its distribution and assess its potential spread in central Tunisia.

Regarding *Cx. pipiens* species, the *pipiens* biotype was predominant in Monastir (56%), whereas in the *hybrid* and *molestus* were most frequently identified in Kairouan (46.1% and 38.4 % respectively). Our results from Kairouan corroborate previous findings regarding the distribution of *Cx. pipiens* forms in this region (Beji et al., 2017). The *pipiens* biotype is typically ornithophilic, preferring to feed on birds, and tends to occupy aboveground habitats, while the *molestus* form is more anthropophilic, feeding on humans, and is adapted to underground or enclosed environments. Hybrids between *pipiens* and *molestus* can exhibit intermediate behaviour, feeding on both birds and humans, which may increase their potential as bridge vectors for zoonotic pathogens such as WNV (Blom et al., 2024), indicating Kairouan as a potential site for WNV transmission to humans.

It’s also important to highlight that *Cx. perexiguus* was found in a high abundance at both sites, which increases the risk of the transmission of WNV, Usutu virus and Sindbis virus in Tunisia (M’Ghirbi et al., 2023).

Monitoring insecticide resistance in both sites revealed the presence of low to moderate frequencies of *kdr* mutations L1014F in *Cx. pipiens* and I1532T in *Ae. albopictus*. The frequency of the *kdr* mutation was notably high in *Cx. pipiens* from Monastir, with a greater proportion of homozygous individuals. This pattern indicates that a sustained selection pressure is acting on the *Cx. pipiens* population in this area. Previous studies have already reported low to high levels of pyrethroid resistance in *Cx. pipiens pipiens* from Tunisia (Daaboub et al., 2008, Ben Cheikh et al., 1998). Our results extend this knowledge by providing the first evidence of the *kdr* mutation in *Cx. pipiens hybrid* and *Cx. pipiens molestus*, and, importantly, the first documentation of homozygous *kdr* alleles in Tunisian *Cx. pipiens*. The presence of *kdr* mutation in *Cx. perexiguus* population, was almost absent (4%). This likely reflects limited or recent exposure of this species to pyrethroid insecticides, due to ecological and behavioural differences from *Cx. pipiens*. Indeed, *Cx. perexiguus* present different breeding sites and biting behaviour compared to *Cx. pipiens*, possibly reducing its contact with insecticide-treated surfaces (Ferraguti et al., 2021).

Regarding OP and CRB resistance, in our sampled populations from both study sites we detected only the G119S mutation, exclusively in the heterozygous state, indicating a low level of resistance to OP and CRB. The G119S allele was found at a low frequency across the three *Cx. pipiens* biotypes, whereas the F290V mutation was absent from all screened specimens. Our findings appear in contradiction with previous study that reported the presence of F290V mutations in Tunisian *Cx. pipiens* populations sampled in 2005 (Ben Cheikh et al., 2009). No resistance associated mutations to DFB, were detected in any of the *Culex* species analysed, suggesting that DFB remains a suitable option for larvicidal applications.

For *Ae. albopictus*, our results indicate the emergence of low-frequency pyrethroid target-site resistance, notably undetected in *Ae. caspius*. The detection of *kdr* III heterozygotes suggests that selection pressure on *Ae. albopictus* is still at an early stage or relatively weak, possibly driven by urban insecticide applications or domestic vector-control practices. Moreover, a moderate CNV of *CCEae3a* and *CCEae6a* associated with OP metabolic resistance was observed in all sampled *Ae. albopictus* specimen. These findings may reflect the introduction of this carboxylesterase’s amplification with the invading population, given that *Ae. albopictus* was only recently detected in Tunisia (2018) and is reported here for the first time in the central region (Bouattour et al., 2019). The upregulation of *CCEae3a* and *CCEae6a* in *Ae. albopictus* in Europe were already described in 2017 and 2020 (Balaska et al., 2020, Grigoraki et al., 2017).

Our findings emphasize the importance of conducting vector surveillance and molecular insecticide resistance profiling in view of supporting vector control in Tunisia. Here we document the spread of *Ae. albopictus* in central Tunisia and provide an updated overview of insecticide resistance in major disease vectors in the country. Our results indicate the suitability of DFB for *Culex* and *Ae. albopictus* control in central Tunisia, emerging pyrethroid resistance in both vector species, and possibly reduced organophosphate efficacy against *Ae. albopictus*. Implementing evidence-based vector control strategies tailored to local ecological settings are key to preventing the spread of insecticide resistance in mosquito vectors in Tunisia and ensuring the continued efficacy of prevention/control interventions.

## Acknowledgements

We acknowledge the support of the European Union’s Marie Skłodowska-Curie postdoctoral fellowship (MSCA) under Grant Agreement No. 101152599. We are very grateful to the Regional Commissariat for Agricultural Development of Kairouan, Tunisia who helped us to accomplish this work.

## Disclosure

The authors declare no conflict of interest.

## Author’s contributions

**LR:** contributed in: the study conception, sampling, molecular analysis, results interpretation, writing of the manuscript and receiver of the MSCA PF, **NK:** molecular analysis and draft writing, **EAF:** draft revision, **LG:** study conception and grant writing, **SB:** molecular analysis, **SL:** specimen collection, **HB:** field work supervision, **NH:** specimen collection, **JV:** study conception, supervision, draft revision and grant writing.

## Notes

### Competing Interest Statement

The authors have declared no competing interest.

